# Unique-region phosphorylation targets LynA for rapid degradation, tuning its expression and signaling in myeloid cells

**DOI:** 10.1101/550053

**Authors:** Ben F. Brian, Adrienne S. Jolicoeur, Candace R. Guerrero, Myra G. Nunez, Zoi E. Sychev, Siv A. Hegre, Pål Sætrom, Nagy Habib, Justin M. Drake, Kathryn L. Schwertfeger, Tanya S. Freedman

## Abstract

The activity of Src-family kinases (SFKs), which phosphorylate immunoreceptor tyrosine-based activation motifs (ITAMs), is a critical factor regulating myeloid-cell activation. We reported previously (Freedman *et al.*, 2015) that the SFK LynA is uniquely susceptible to rapid ubiquitin-mediated degradation in macrophages, functioning as a rheostat regulating signaling. We now report the mechanism by which LynA is preferentially targeted for degradation and how cell specificity is built into the LynA rheostat. Using genetic, biochemical, and quantitative phosphopeptide analyses, we found that the E3 ubiquitin ligase c-Cbl preferentially targets LynA via a phosphorylated tyrosine (Y32) in its unique region. This distinct mode of c-Cbl recognition depresses steady-state expression of LynA in macrophages derived from mice. Mast cells, however, express little c-Cbl and have correspondingly high LynA. Upon activation, mast-cell LynA is not rapidly degraded, and SFK-mediated signaling is amplified relative to macrophages. Cell-specific c-Cbl expression thus builds cell specificity into the LynA checkpoint.

## Introduction

Phosphorylation of immunoreceptor tyrosine-based activation motifs (ITAMs) by Src-family kinases (SFKs) is the first step in the activation of an innate immune response during a pathogen encounter. Initiation of cell-activating signaling typically occurs within clusters of ITAM-coupled receptors such as FcγR (1) or the hemi-ITAM Dectin-1 (2, 3), nucleated by highly multivalent binding to immunoglobulin-G-decorated pathogens or fungal cell-wall β-glucans respectively. Together, SFKs and phosphorylated ITAMs trigger activation of the tyrosine kinase Syk (4). The SFKs and Syk then drive activation of membrane-proximal signaling through adaptor proteins, cytoskeleton-modulating proteins such as the FAK/Pyk2 kinases and Paxillin, and Tec kinases, which activate second-messenger pathways via phosphoinositide 3-kinase and phosphoinositide phospholipases Cγ (PI3K and PLCγ1/2 respectively). Ultimately, these early events lead to downstream signaling through Erk1/2 and other pathways (4). As the byproducts of activated macrophages can be toxic (*e.g.* release of reactive oxygen species) and drive inflammation (*e.g.* release of tumor necrosis factor α), the responsiveness of innate immune cells must be tightly regulated (5–8).

Multiple mechanisms work together to tune the responsiveness of macrophages and other myeloid cells, including negative regulation by the phosphatases CD45 and CD148 (5, 9, 10), cytoskeletal barriers to diffusion (11), signaling via immunoreceptor tyrosine inhibitory motifs (ITIMs) (12) and inhibitory ITAMs (13, 14), and degradation and mislocalization of signaling molecules targeted for polyubiquitination by ubiquitin ligases (15, 16). The SFKs, which in myeloid cells typically include Fgr, Fyn, and two splice forms each of Hck and Lyn, may also have positive and negative functions (12, 17–19). Layered onto the traditional positive- and negative-regulatory roles of the SFKs, we have shown that activated LynA (the longer of the two Lyn splice forms) is rapidly and specifically targeted for polyubiquitination and degradation, forming the basis of a signaling checkpoint that blocks spurious macrophage activation (20). The mechanism preferentially targeting LynA for polyubiquitination and the molecular basis of this selectivity have not been elucidated previously.

Unlike Dectin-1 and FcγR signaling in macrophages, which is typically triggered in the context of pathogen-induced µm-scale clusters of receptors (5, 9, 20), mast-cell FcεR signaling has a low threshold for activation-- small or even monovalent antigen-IgE complexes are sufficient to induce a signaling response (21, 22). Like macrophages, mast cells express LynA, and this disparity in receptor sensitivity has not previously been explained.

This paper describes the mechanism by which active LynA is selectively recognized and rapidly degraded, tuning its activation kinetics and its steady-state protein levels in a cell-specific manner. To reveal the requirements for LynA degradation, we synchronized receptor-independent SFK activation using the designer inhibitor 3-IB-PP1 (23–25), which specifically inhibits a variant of the SFK-inhibitory kinase Csk (Csk^AS^) (20, 25, 26). Csk is responsible for phosphorylating a key inhibitory tyrosine in the C-terminus of all the SFKs. Inhibiting Csk^AS^ leads to rapid and robust SFK activation due to the loss of the dynamic equilibrium between Csk and the phosphatases CD45 and CD148 (27, 28). We showed previously that SFK activation induced via 3-IB-PP1 treatment leads to the phosphorylation of the E3 ubiquitin ligase c-Cbl and the preferential polyubiquitination and degradation of the SFK LynA, but it was not clear whether these two events were linked. Using knockdown, overexpression, and genetic models, we now demonstrate that c-Cbl, but not its homolog Cbl-b, controls the steady-state expression of LynA and mediates its rapid degradation upon activation in macrophages. LynA is targeted by c-Cbl via a phosphorylation site, Tyr32 within its unique region, a mode of interaction distinct from the slower-phase action of c-Cbl and other E3 ligases on LynB and the other SFKs. Finally, we have discovered that the LynA checkpoint is cell specific. In mast cells, which express very little c-Cbl but abundant Cbl-b, LynA is not rapidly degraded upon activation, helping to explain why mast cells have a lower activation threshold than macrophages.

## Results

### c-Cbl mediates steady-state and activation-induced degradation of LynA in macrophages

We reported previously that activated LynA is rapidly polyubiquitinated and degraded in Csk^AS^ macrophages treated with the Csk^AS^ inhibitor (SFK activator) 3-IB-PP1 (20). To identify the responsible E3 ubiquitin ligase, we tested the effects of knocking down the expression of candidate ligases using small interfering (si)RNA. Csk^AS^ bone-marrow-derived macrophages (BMDMs) were transfected with non-targeting control RNA or with siRNA constructs targeting c-Cbl or Cbl-b, known modulators of ITAM signaling in both adaptive and innate immune cells (12, 15, 29). Resting Csk^AS^ BMDMs were treated with 3-IB-PP1 for up to 5 min to activate the SFKs and initiate bulk degradation of LynA. Immunoblots of whole-cell lysates showed efficient and specific knockdown **(Figure 1A)** of c-Cbl protein (70% reduced) and Cbl-b protein (60% reduced) **(Figure 1B)**.

**Figure 1.**
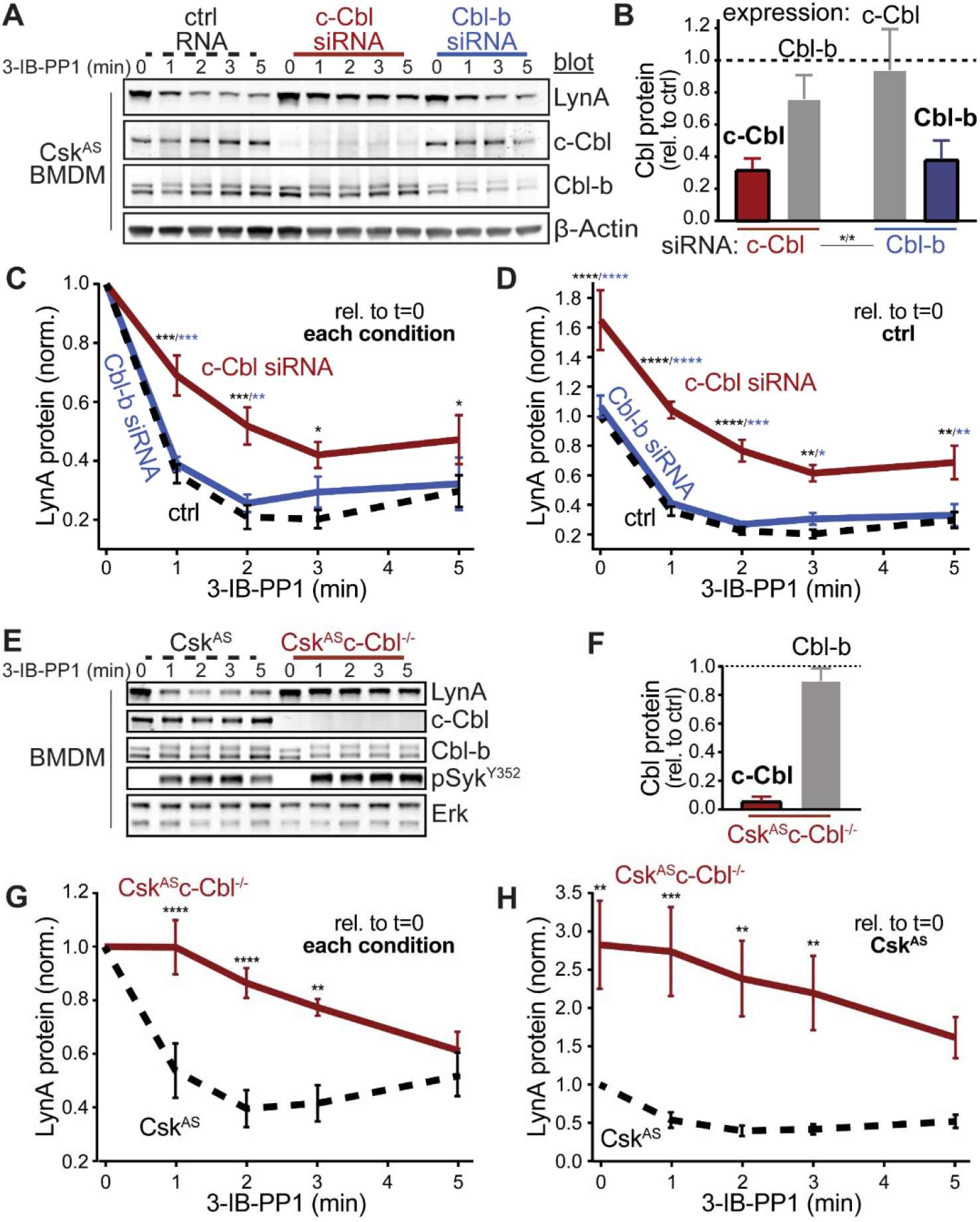
Loss of c-Cbl expression in macrophages leads to steady-state upregulation and delayed degradation of LynA protein. **(A)** Immunoblots showing LynA and Cbl protein content of Csk^AS^ BMDMs transfected with non-targeting control RNA (ctrl, black) or siRNA sequences targeting c-Cbl (red) or Cbl-b (blue). After overnight incubation, cells were treated with 3-IB-PP1 for the indicated times; β-actin is shown as a loading control. **(B-D)** Densitometry quantification of relative Cbl and LynA protein levels, corrected for total protein content. Error bars reflect standard error of the mean (SEM), n=3. **(B)** Quantification of c-Cbl and Cbl-b relative to ctrl expression of each (dotted line). Significant differences (Sig.) from two-way ANOVA with Sidak’s multiple comparison test (asterisks): [c-Cbl expression: c-Cbl vs. Cbl-b siRNA samples], [Cbl-b expression: c-Cbl vs. Cbl-b siRNA samples]; * P=0.0217-0.0473. **(C-D)** Quantification of LynA relative to t=0 within each siRNA condition **(C)** or the steady-state level in the ctrl **(D)**. Sig. from two-way ANOVA with Tukey’s multiple comparison test: [c-Cbl vs. ctrl, black asterisks], [c-Cbl vs. Cbl-b, blue asterisks]; **** P<0.0001, *** P=0.0001-0.0005, ** P=0.0015-0.0058, * P=0.0105-0.0451. No significant differences (ns) between other pairs. **(E)** Immunoblots showing LynA and Cbl protein content of Csk^AS^ and Csk^AS^c-Cbl^−/−^ BMDMs treated with 3-IB-PP1; Erk1/2 is shown as a loading control. **(F-H)** Quantification of relative Cbl and LynA protein levels, corrected for total protein content. SEM, n=5. **(F)** Quantification of c-Cbl and Cbl-b in Csk^AS^c-Cbl^−/−^ BMDMs relative to expression levels in Csk^AS^ BMDMs (dotted line). **(G-H)** Quantification of LynA relative to t=0 for each genotype **(G)** or the steady-state level in Csk^AS^ BMDMs **(H)**. Sig. from two-way ANOVA with Tukey’s multiple comparison test: [Csk^AS^ vs. Csk^AS^c-Cbl^−/−^, asterisks]; **** P<0.0001, *** P=0.0004, ** P=0.0014-0.0048. Paradoxical loss of Erk signaling in c-Cbl-deficient cells is shown in Figure 1–figure supplement 1. Refer to Figure 1–Source Data 1.

Knockdown of c-Cbl led to a delay in the degradation of activated LynA, with 1 min 3-IB-PP1 treatment reducing LynA by 30% in c-Cbl knockdown BMDMs compared to 61% and 64% in Cbl-b knockdown and control BMDMs, respectively **(Figure 1C)**. c-Cbl knockdown also increased the level of steady-state LynA by 60% **(Figure 1D)**. The impaired degradation and increased steady-state LynA expression together resulted in pronounced elevations in LynA protein in the first minutes of 3-IB-PP1 treatment; after 1 min of 3-IB-PP1 treatment, LynA was elevated 2.9-fold in cCbl-deficient BMDMs relative to control. Although it has been shown previously that c-Cbl and Cbl-b can have redundant functions (30, 31), we were unable to detect any role for Cbl-b in the steady-state or activation-induced degradation of LynA.

To test whether the residual LynA degradation was due to incomplete knockdown, we also generated *Csk^AS^Cbl^−/−^* mice to test LynA degradation in BMDMs lacking any c-Cbl expression. As with siRNA knockdown, total c-Cbl knockout impaired LynA degradation in BMDMs **(Figure 1E)**; Cbl-b expression in Csk^AS^c-Cbl^−/−^ BMDMs remained comparable to wild-type levels **(Figure 1F)**. Complete loss of c-Cbl caused a more profound impairment of LynA degradation during the first few minutes of 3-IB-PP1 treatment **(Figure 1G)** a dramatic increase in steady-state LynA protein (up to 6-fold higher than in c-Cbl^+/+^ cells, **Figure 1H**). As in the knockdown experiments, LynA degradation in c-Cbl^−/−^ BMDMs was not fully eliminated but occurred on a much slower timescale, similar to the degradation of the other SFKs (20). From this observation we conclude that c-Cbl is solely responsible for the rapid-phase degradation unique to LynA, whereas other E3 ubiquitin ligases can complement c-Cbl to mediate slower-phase SFK degradation. Although activating Syk phosphorylation was increased in Csk^AS^c-Cbl^−/−^ BMDMs relative to Csk^AS^ BMDMs treated with 3-IB-PP1 **(Figure 1E)**, Erk phosphorylation after SFK activation was paradoxically impaired **(Figure 1–figure supplement 1)**, likely due to the loss of the adaptor function of c-Cbl in PI3K signaling (32, 33).

### A tyrosine residue in the unique-region insert of LynA is required for its rapid degradation

Susceptibility to rapid, activation-induced degradation differentiates LynA from the splice variant LynB and the other macrophage SFKs. Hck, Fgr, LynB, and Fyn (17, 19), are degraded >10-fold more slowly than LynA during 3-IB-PP1 treatment (20) **(Figure 2A)**. Although not expressed in macrophages, Lck is also degraded more slowly than LynA **(Figure 2B)**.

**Figure 2.**
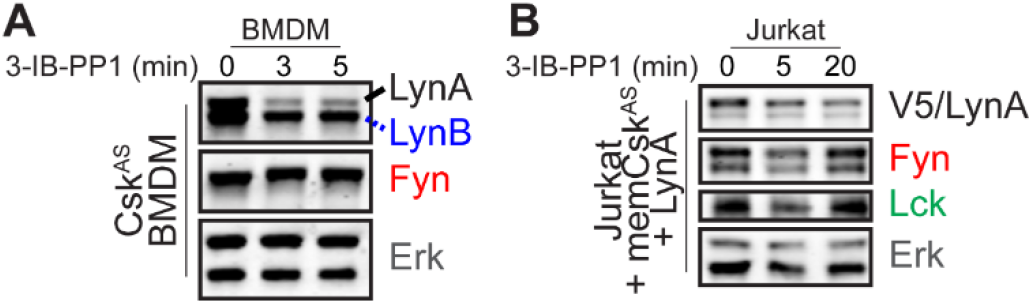
Fyn and Lck are not rapidly degraded upon activation. Immunoblots and quantification of protein levels of Fyn, Lck, and LynB in 3-IB-PP1-treated Csk^AS^ BMDMs **(A)** and Jurkat cells **(B)**. Total Erk1/2 protein (Erk) is a loading control.

As we have been unable to overexpress LynA in BMDMs by transfection, by Amaxa nucleofection, or by lentiviral transduction, we expressed LynA ectopically in Jurkat T cells by transient transfection along with a Cbl construct and a membrane-localized construct of Csk^AS^ (memCsk^AS^) (25), which sensitizes Jurkat cells to 3-IB-PP1 for synchronized SFK activation. As a proof of principle for this model system, we transfected Jurkat cells with wild-type LynA, memCsk^AS^, and empty vector (ctrl), c-Cbl, or Cbl-b plasmid DNA **(Figure 3A)**, which increased c-Cbl expression 3-fold and Cbl-b 25-fold relative to endogenous levels **(Figure 3B)**. Overexpressing one Cbl family member did not affect the abundance of the other. As predicted, c-Cbl overexpression in Jurkat cells increased the rate of LynA degradation (30% depleted after 5 min 3-IB-PP1 treatment in c-Cbl-overexpressing Jurkat cells vs. 10% and 0% in ctrl and Cbl-b-overexpressing cells, respectively) **(Figure 3C)**. This observation is consistent with our siRNA experiments in BMDMs, suggesting that Jurkat cells, which do not normally express Lyn, are capable of supporting rapid, c-Cbl-mediated degradation of LynA.

**Figure 3.**
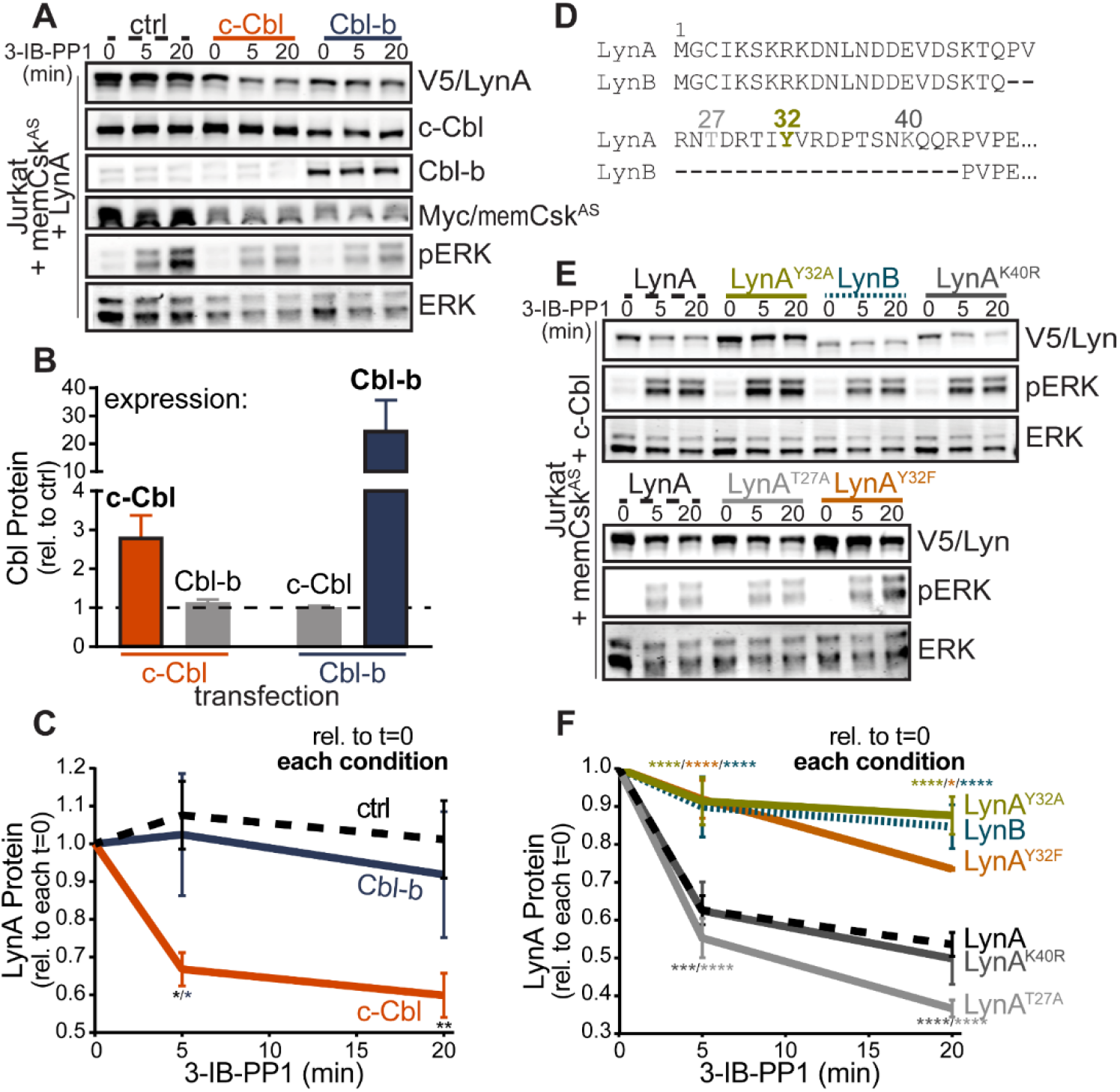
A unique tyrosine residue is required for activation-induced degradation of LynA. **(A)** Immunoblots of Jurkat cells cotransfected with V5-tagged LynA, Myc-tagged memCsk^AS^, and either empty vector (ctrl, black dotted), c-Cbl (orange), or Cbl-b (blue) and treated with 3-IB-PP1. Activating phosphorylation (Thr202/Tyr204) of ERK1/2 (pERK) shows activating signaling, and total ERK1/2 (ERK) shows lane loading. **(B-C)** Quantification of relative Cbl and LynA protein, corrected for total protein content. SEM, n=4. **(B)** Quantification of overexpressed c-Cbl and Cbl-b relative to endogenous (ctrl) levels (dotted line). **(C)** Quantification of LynA protein during 3-IB-PP1 treatment relative to the steady-state level. Sig. from two-way ANOVA with Tukey’s multiple comparison test: [c-Cbl vs. ctrl, black asterisks], [c-Cbl vs. Cbl-b, blue asterisks]; ** P=0.0092, * P=0.0270-0.0102. Other pairs ns. **(D)** N-terminal amino-acid sequences of mouse LynA and LynB (34, 35), including the 21 amino-acid unique insert of LynA. Tyrosine 32 (olive), threonine 27 (light gray), and lysine 40 (dark gray) are labeled. **(E)** Immunoblots of 3-IB-PP1-treated Jurkat cells cotransfected with c-Cbl, Myc-tagged memCsk^AS^, and V5-tagged Lyn constructs, including wild-type LynA (black, dashed), LynA^Y32A^ (olive), LynB (teal, dotted), LynA^K40R^ (dark gray), LynA^T27A^ (light gray), or LynA^Y32F^ (orange). (F) Quantification of LynA protein levels, corrected for total protein, relative to the steady-state level. SEM, n=7 for LynA, n=4 for LynA^Y32A^ and LynB; n=3 for LynA^K40R^, LynA^T27A^, and LynA^Y32F^. Sig. from two-way ANOVA with Tukey‘s multiple comparison test: [LynA^Y32A^ vs. LynA, olive asterisks], [LynB vs. LynA, teal asterisks], [LynA^Y32F^ vs. LynA, orange asterisks], [LynA^K40R^ vs. LynA^Y32A^, dark gray asterisks], [LynA^T27S^ vs. LynA^Y32A^, light gray asterisks]; **** P<0.0001, *** P=0.0006, * P=0.0166. ns: [LynA^T27S^ and LynA^K40R^ vs. each other and LynA] and [LynA^Y32A^ and LynA^Y32F^ vs. each other and LynB]. Note: the error bar for LynA^Y32F^ is smaller than the line width. Relative expression levels of LynA variants and LynB are shown in Figure 3–figure supplement 1. Refer to Figure 3–Source Data 1.

LynB, which lacks a 21 amino-acid insert found in the unique region of LynA **(Figure 3D)**, was less efficiently degraded in c-Cbl-transfected Jurkat cells **(Figure 3E)**, depleted by 10% (LynB) vs. 40% (LynA) from baseline after 5 min 3-IB-PP1 **(Figure 3F)**. These data mirror our findings in Csk^AS^ BMDMs (20) and confirm that the unique sensitivity of LynA to rapid degradation is preserved in the Jurkat model. We hypothesized that the targeting site in LynA must lie within the unique-region insert (residues 23-43) lacking in LynB (34, 35), which contains a tyrosine phosphorylation site (Tyr32) (36), a predicted ubiquitination site (Lys40) (UbPred, 37), and a predicted threonine phosphorylation site (Thr27) (NetPhos 3.1, 38) **(Figure 3D)**.

Substituting LynA Tyr32 with either alanine or phenylalanine completely blocked its rapid degradation **(Figure 3E)**, reducing the depletion after 5 min 3-IB-PP1 from 40% (wt) to 10% (LynA^Y32A^) or 8% (LynA^Y32F^) from baseline, levels indistinguishable from LynB **(Figure 3F)**. In contrast, mutation of the predicted ubiquitination (LynA^K40R^) or threonine phosphorylation (LynA^T27A^) sites **(Figure 3E)** did not impair LynA degradation **(Figure 3F)**.

As in BMDMs, the efficiency of Lyn degradation also affects its steady-state expression in Jurkat cells. LynB is 2- and LynA^Y32A^ 3-fold more highly expressed than wild-type LynA, LynA^K40R^, or LynA^T27A^ in resting cells, resulting in 3- to 4-fold more protein remaining after 20 min treatment with 3-IB-PP1 **(Figure 3–figure supplement 1)**. Overall, we conclude that the unique tyrosine residue in the LynA insert, a known site of tyrosine phosphorylation (36), is the recognition site for rapid, c-Cbl-mediated degradation, and that this mechanism determines the half-life of activated LynA protein within an ITAM signaling complex and tunes the steady-state expression of LynA protein. This suggests that c-Cbl-mediated regulation of steady-state LynA protein levels occurs as a direct effect of c-Cbl protein activity rather than merely a transcriptional feedback effect.

### LynA kinase activity is required for its rapid degradation

SFK activation is a requirement for rapid degradation of LynA protein in BMDMs (20), but whether this is direct effect of the LynA kinase itself or initiated by feedback from downstream signaling proteins has been unclear. To investigate the role of LynA kinase activity in its degradation, we tested two functionally impaired LynA constructs. First, we introduced a lysine residue in place of an activation-loop threonine (T410K, c-Src residue 429 (39)) to disrupt substrate recognition (40). Second, we introduced a phenylalanine residue in place of the activation-loop tyrosine, a known site of interaction between the phosphotyrosine binding (PTB) domain of c-Cbl and Src (41) and the key autophosphorylation site that stabilizes the active state (27, 42) (Y397F, c-Src residue 416).

In untransfected Jurkat cells ERK1/2 is phosphorylated downstream of the endogenous SFKs Lck (43, 44) and Fyn (45), which are activated following treatment with 3-IB-PP1 (25). Since the endogenous kinases are not disrupted by substitutions in transfected LynA, ERK1/2 continues to be efficiently phosphorylated in cells transfected with LynA T410K or Y397F **(Figure 4A)**. In JCaM1.6 cells, an Lck-deficient Jurkat strain (43, 46), ERK1/2 phosphorylation depended on transfected LynA and reflected the functional impairment of LynA T410K and Y397F **(Figure 4B)**. As reported previously, Fyn activity in the absence of Lck cannot mediate ERK1/2 phosphorylation (43).

**Figure 4.**
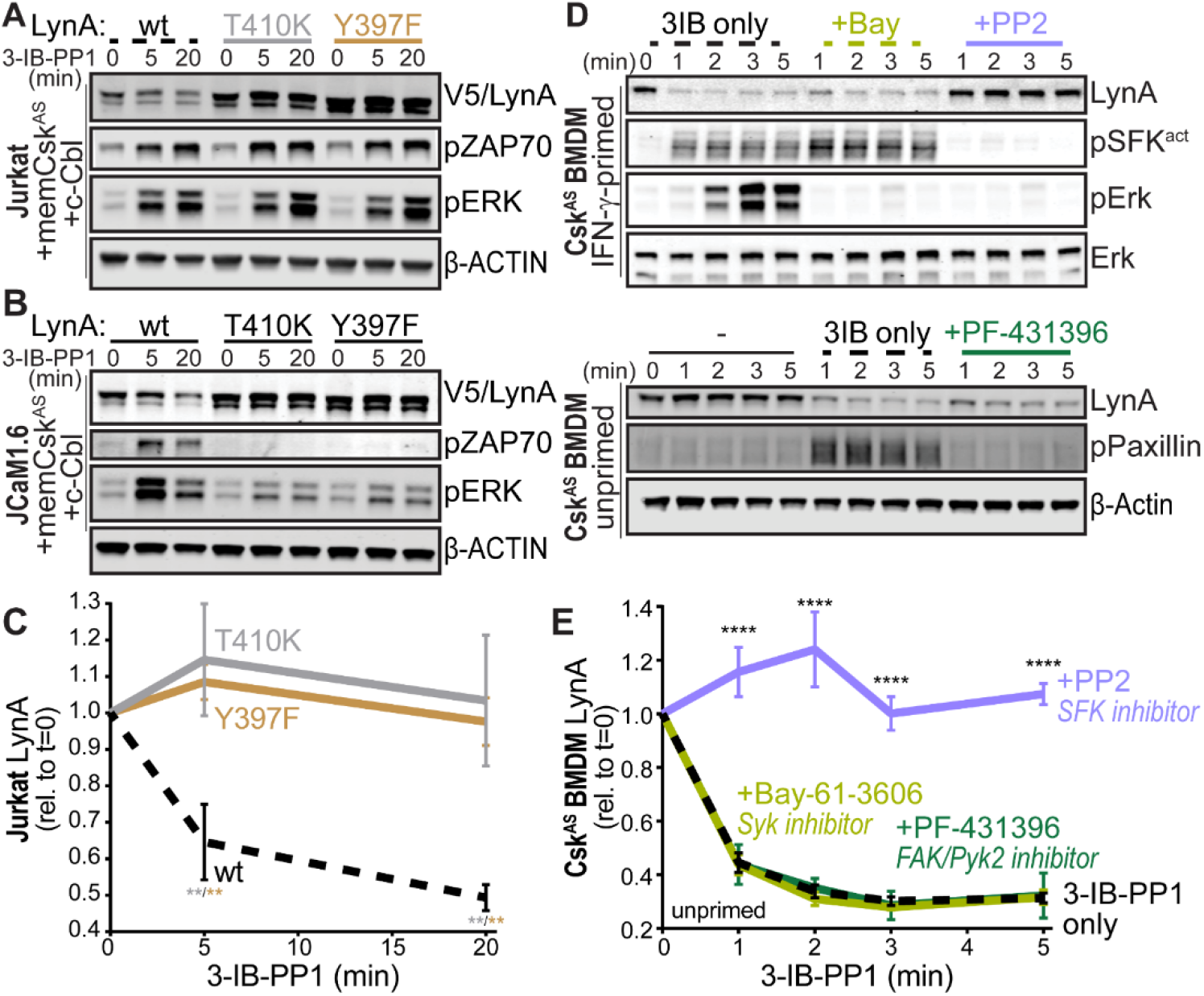
SFK catalytic activity is required for rapid degradation of LynA. **(A)** Immunoblots showing ectopic expression of wild-type (wt, black dotted), substrate-binding-impaired (T410K, gray), or activation-loop-mutated (Y397F, orange) V5-tagged LynA protein coexpressed in Jurkat cells with memCsk^AS^ and c-Cbl and treated with 3-IB-PP1. pERK is shown as a control for 3-IB-PP1 treatment. β-actin reflects lane loading. **(B)** Overexpression of wt and variants of LynA in JCaM1.6 cells. ZAP70 (Y319) and ERK are phosphorylated as a consequence of SFK activation. **(C)** Quantification of LynA protein in transfected Jurkat cells during 3-IB-PP1 treatment, corrected for total protein, relative to the steady-state level for each variant. SEM, n=3. Sig. from two-way ANOVA with Tukey’s multiple comparison test: [T410K vs. wt, gray asterisks], [Y397F vs. wt, orange asterisks]; ** P=0.0015-0.0083. Other pairs ns. **(D)** Immunoblots showing expression of endogenous LynA protein in Csk^AS^ BMDMs treated with 3-IB-PP1 (3IB only, black dotted) or with 3-IB-PP1 plus the Syk inhibitor Bay-61-3606 (Bay, light olive), the SFK inhibitor PP2 (light purple), or the FAK/Pyk2 inhibitor PF-431396 (green). Phosphorylated SFK activation loop Tyr416 (pSFK^act^), Erk1/2 (pErk), and Paxillin Tyr118 (pPaxillin) are shown as controls for inhibitor function; β-actin and Erk are shown as loading controls. (E) Quantification of LynA protein during 3-IB-PP1 treatment, corrected for total protein, relative to the steady-state level. SEM, n=3. Sig. from two-way ANOVA with Tukey’s multiple comparison test: [+PP2vs. 3IB only, asterisks]; **** P<0.0001. Other pairs ns. Refer to Figure 4–Source Data 1.

Functional LynA kinase was required for c-Cbl-mediated degradation in Jurkat cells. While wild-type LynA protein was 50% depleted after 20 min of 3-IB-PP1 treatment, degradation of either LynA T410K or Y397F was undetectable **(Figure 4C)**. The dependence on LynA kinase function was evident in both Lck- and Fyn-expressing Jurkat cells **(Figure 4A)** and Lck-deficient JCaM1.6 cells **(Figure 4B)**, indicating that, in this system, the other activated SFKs could not target LynA in trans to induce c-Cbl-mediated degradation. We do not yet know whether the unique insert of LynA is a substrate for the other macrophage SFKs, but the peptide containing Tyr32 is a predicted SFK substrate (NetPhos 3.1, 38).

In macrophages and other hematopoietic cells the SFKs are the first kinases to be activated upon antigen receptor engagement, phosphorylating intracellular ITAMs, ITAM-associated Syk or Zap70, and integrin-associated FAK or Pyk2 (4, 47). We have reported that Syk and FAK are phosphorylated on activating tyrosine residues in Csk^AS^ BMDMs treated with 3-IB-PP1 (20). To test whether these downstream kinases might participate in negative feedback leading to LynA degradation, we treated Csk^AS^ macrophages with competitive inhibitors of Syk (48), FAK/Pyk2 (49), or the SFKs themselves (50) in combination with 3-IB-PP1. As controls for on-target effects, we demonstrated that Erk1/2 and SFK activation-loop phosphorylation was blocked in samples cotreated with the pan-SFK inhibitor PP2, that Erk1/2 but not SFK activation-loop phosphorylation was blocked in samples cotreated with the Syk inhibitor BAY-61-3606, and that Paxillin phosphorylation was blocked in samples cotreated with the FAK/Pyk2 inhibitor PF-431396 **(Figure 4D)**. As with genetically impairing LynA kinase function, inhibiting SFK activity with PP2 abrogated LynA degradation **(Figure 4D-E)**. In contrast, LynA degradation in BMDMs was unaffected by cotreatment with BAY-61-3606 or PF-431396. Together these data suggest that Syk and FAK/Pyk2 (and any off-target effects of BAY-61-3606 and PF-431396) do not participate in a larger negative-feedback process, suggesting that LynA might target itself directly through Tyr32 in cis or trans. Interestingly, since the T-cell SFKs Lck and Fyn cannot effect LynA degradation in trans, this feedback process could be limited to a kinase effect in cis or to another macrophage-expressed Src family member. This tight feedback circuit could explain the strikingly fast kinetics of LynA degradation.

### Differential Cbl expression tunes LynA protein levels and signaling responses in macrophages and mast cells

Analysis of data from the Immunological Genome Project shows that expression levels of Lyn and c-Cbl RNA are differentially regulated across different cell types. Mast cells, for instance, express very low levels of c-Cbl mRNA **(Figure 5)** (51, 52), which corresponds to a low level of c-Cbl protein expression (53). Cbl-b mRNA, in contrast, is abundant in mast cells.

**Figure 5.**
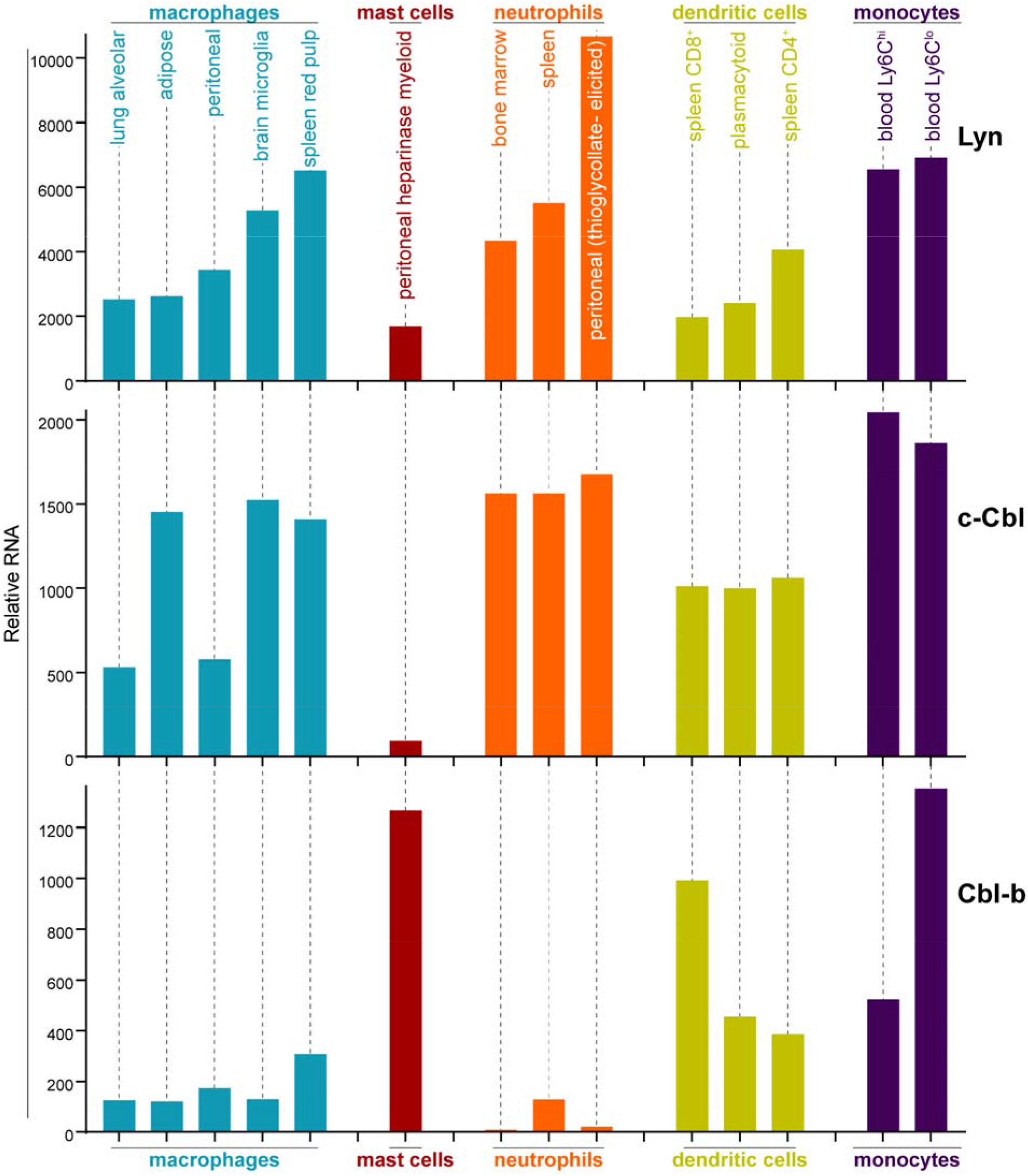
LynA, c-Cbl, and Cbl-b mRNAs are regulated differentially in myeloid cells. Mouse RNAseq data were derived from the Immunological Genome Project (ImmGen, http://rstats.immgen.org/Skyline/skyline.html, 51, 52). RNA levels of Lyn, c-Cbl, and Cbl-b are shown for macrophages, mast cells, neutrophils, dendritic cells, and blood monocytes. Refer to Figure 5–Source Data 1.

As in macrophages, Lyn is a key regulator of mast-cell ITAM signaling (54), but the effect of disparate Cbl-family expression on LynA in the two cell types remains unknown. We generated bone-marrow-derived mast cells from Csk^AS^ mice (hereafter referred to as “mast cells”) and performed surface marker analysis as we had done already for BMDMs (20). Csk^AS^ mast cells were homogenous in their expression of traditional mast-cell markers FcεR1α and c-Kit (CD117) (55) **(Figure 6A)**.

**Figure 6.**
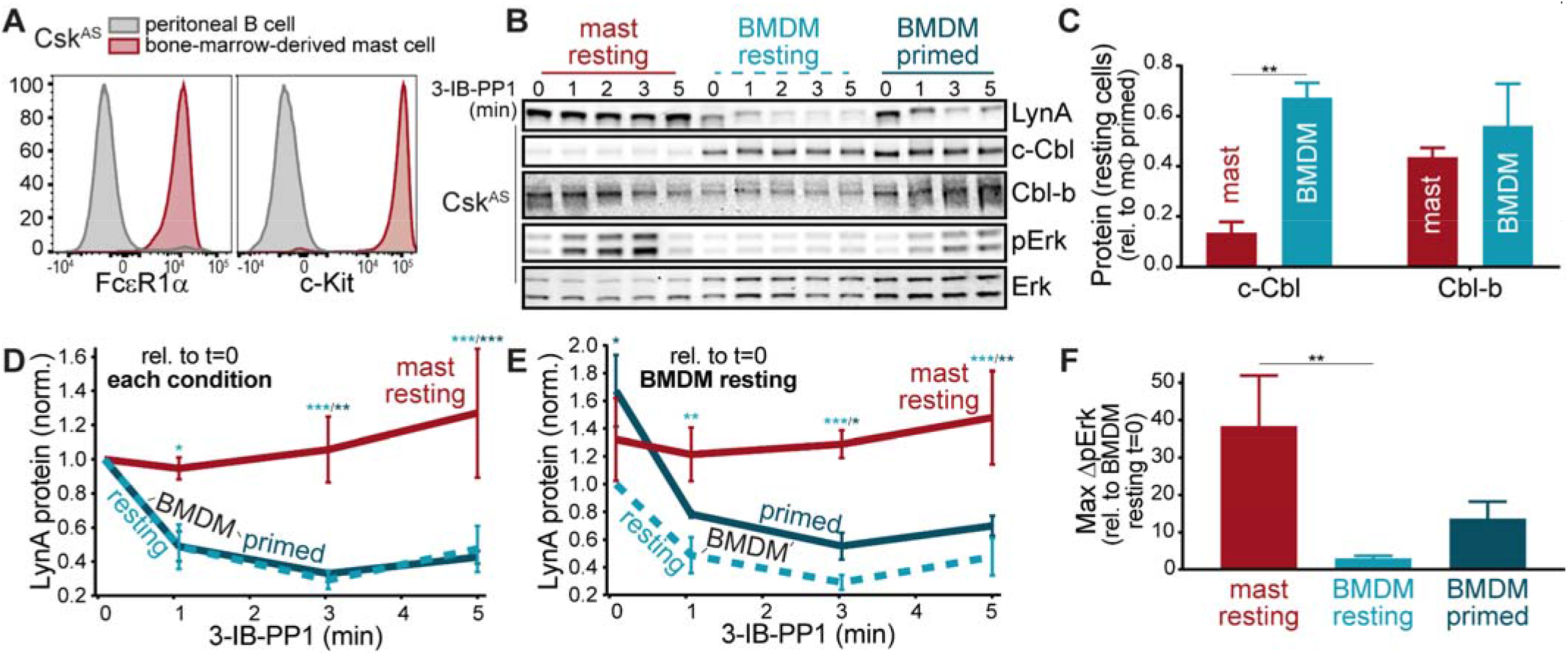
c-Cbl expression correlates with LynA protein levels and pathway activation in macrophages and mast cells. **(A)** Flow cytometry plots showing surface expression of the mast-cell markers FcεR1α and c-Kit (CD117) in Csk^AS^ bone-marrow-derived mast cells compared to a peritoneal B-cell control. **(B)** Immunoblots of Csk^AS^ bone-marrow-derived mast cells (mast resting, red) and BMDMs after resting (light teal) or IFN-γ priming (dark teal). LynA, c-Cbl, Cbl-b, and pErk1/2 are shown; total Erk1/2 reflects lane loading. **(C)** Quantification of c-Cbl and Cbl-b protein in mast cells and macrophages, corrected for total protein, relative to the level in primed macrophages (mΦ). SEM, n=3. Sig. from one-way ANOVA with Sidak multiple comparison test: [mast vs. BMDM resting, asterisk]; ** P= 0.0035. Others pairs ns. **(D-E)** Quantification of LynA, corrected for total protein, relative to the steady-state level in each cell condition **(D)** or in resting BMDMs at t=0 **(E)**. SEM, n=3 for mast cells and primed BMDMs, 4 for resting BMDMs. Sig. from two-way ANOVA with Tukey’s multiple comparison test: [mast vs. BMDM resting, light teal asterisks], [mast vs. BMDM primed, dark teal asterisks]; *** P=0.0004-0.0008, ** P=0.0025-0.0088, * P=0.0134-0.0466. Other pairs ns. **(F)** Quantification of the maximum increase in pErk1/2 over the first 5 min of 3-IB-PP1 treatment, corrected for total Erk1/2 protein, relative to the level in resting BMDMs at t=0. SEM, n=3 for mast cells, 5 for resting BMDMs, and 4 for primed BMDMs. Sig. from one-way ANOVA with Tukey’s multiple comparison test: [mast vs. BMDM resting, light teal asterisks]; ** P= 0.0085. Others pairs ns. Refer to Figure 6–Source Data 1.

We then used immunoblotting to compare the expression of Cbl and LynA protein in Csk^AS^ mast cells and BMDMs **(Figure 6B)** and found that mast cells express very little c-Cbl (20% of the level in resting macrophages) and relatively high levels of Cbl-b (80% of the level in resting macrophages) **(Figure 6C)**. Strikingly, low c-Cbl expression was accompanied by a complete resistance of LynA to degradation in the first 5 min of 3-IB-PP1 treatment **(Figure 6D)** and an increase in steady-state expression of LynA (30% higher than in resting BMDMs) **(Figure 6E)**, resulting in 3-4x more LynA protein in mast cells than in resting BMDMs and 2x more than in primed BMDMs between 3 and 5 min of 3-IB-PP1 treatment. Mast cells also had a stronger signaling response to 3-IB-PP1. Although the kinetics of Erk phosphorylation in mast cells varied among experiments, the fold increase in pErk1/2 in the first 5 min of 3-IB-PP1 treatment was consistently higher than in macrophages, even primed BMDMs (40x pErk increase in mast cells compared with 14x in primed BMDMs and no significant increase in resting BMDMs) **(Figure 6F)**. We therefore conclude that c-Cbl expression tunes the steady-state expression of LynA protein and the cell’s ability to target LynA for rapid degradation upon activation. Since mast cells are less dependent on c-Cbl for positive signaling, long-lived LynA potentiates a strong response to 3-IB-PP1. Thus mast cells bypass the LynA signaling checkpoint by maintaining low levels of c-Cbl and therefore high levels of activated LynA.

## Discussion

Both c-Cbl and Cbl-b negatively regulate ITAM signaling, recognizing activated signaling proteins via PTB binding to activating phosphotyrosine sites. Cbl-mediated polyubiquitination leads to protein degradation via the lysosome and proteasome (56). Cbl-family ligases must in turn be tightly regulated to prevent hyperresponsive or aberrant signaling while maintaining proper pathogen clearance (57). Although highly homologous, c-Cbl and Cbl-b target different subsets of signaling proteins for degradation (58, 59), with c-Cbl targeting the SFKs, including Lyn (60, 61), and Cbl-b interacting with ITAM-coupled receptors and other kinases (31, 53, 57). c-Cbl is thought to interact with active SFKs through the SFK activation-loop phosphotyrosine residue, which binds the c-Cbl PTB domain (62), and through the SFK SH3 domain, which binds the c-Cbl PXXP motifs (63). In this paper we report a new mode of recognition specific to LynA. Although LynA and LynB have identical activation kinetics and activation-loop and SH3-domain sequences, LynB is degraded 10-fold more slowly in macrophages treated with 3-IB-PP1 (20). This rapid degradation profile is unique among the SFKs we have tested (LynB, HckA, HckB, Fgr (20), and Fyn in BMDMs and Lck and Fyn in Jurkat cells). By exploiting the similarity between LynA and LynB, we conclude that, while activation-loop phosphorylation and SH3-domain interactions mediate slower-phase targeting by c-Cbl and other E3 ligases, rapid degradation of LynA is triggered by an interaction unique to its insert region.

A single tyrosine residue (Tyr32) peculiar to its unique region marks activated LynA for rapid degradation. This phenomenon is easily observed upon bulk SFK activation by 3-IB-PP1 but also affects the steady-state level of LynA protein, likely due to the degradation of basally active LynA over time. Phosphorylation of LynA Tyr32 has been observed in neuroblastoma cell lines upon receptor-tyrosine-kinase activation (64). Further, in epidermoid carcinoma (A431) cells and breast tumor samples LynA Tyr32 is phosphorylated by the epidermal growth factor receptor (EGFR), allowing it to phosphorylate MCM-7 and stimulate cell proliferation (36). In macrophages this potentially positive-regulatory function of Tyr32 paradoxically causes LynA to trigger its own c-Cbl-mediated degradation, underlying its function as a rheostat in the LynA checkpoint. We can speculate about two possible mechanisms. First, by analogy to a similarly situated but non-homologous transautophosphorylation site in Hck (Tyr^29^) (65), LynA phospho-Tyr32 may be directly activating. The unique regions of LynA and LynB make distinct SH3-domain contacts (66), and phospho-Tyr32 could alter these interactions. Hyperactivated LynA could then efficiently phosphorylate and activate nearby or pre-complexed c-Cbl, triggering its own degradation more aggressively than do the other SFKs. Alternatively, phospho-Tyr32 could directly or indirectly serve as a docking site for c-Cbl, constituting a unique interaction site to efficiently recruit its degradation machinery. Bolstering this hypothesis is a published study showing that the interactome of LynA^Y32F^ transfected into mammary adenocarcinoma (MDA-MB-231) cells more closely resembles the interactome of wild-type LynA than LynB, but some LynA interactions are lost (67). It is also possible that LynA Tyr32 is not phosphorylated, but is required for an allosteric intramolecular interaction or docking of interacting proteins.

We have shown that SFK activity is required for LynA degradation, and LynA Tyr32 is a predicted substrate for Hck (NetPhos 3.1, 38) in addition to EGFR. It is therefore tempting to speculate that LynA Tyr32 is a site of autophosphorylation, since the SFKs Lck and Fyn cannot mediate the degradation of kinase-impaired LynA in trans. On the other hand, we have shown that LynA and LynB are activation-loop phosphorylated 20x more quickly than Hck and Fgr in response to 3-IB-PP1 (20), and it is possible that this delay could function as a timer, allowing a rapid burst of LynA activation with a short half-life, either globally (in the case of 3-IB-PP1 treatment) or at the single-molecule scale within a phagocytic synapse.

LynA and LynB have been reported to interact with different subsets of proteins in mast cells and in triple-negative breast cancer cells, with LynA signaling via ITAM (18), cytoskeletal, and proliferative (67) pathways and LynB initiating negative feedback via ITIMs and phosphatases (18). In mammary epithelial cells the ratio of LynA and LynB is actively regulated by Epithelial Splicing Regulatory Protein 1 (ESRP1), with LynA-upregulated tumors having the more invasive phenotype (67). The positive-regulatory roles reported for LynA in mast and tumor cells complements our own observations that LynA degradation can block macrophage signaling through the Erk, Akt, and NFAT pathways, which cannot be complemented by active LynB (20).

Aberrant activation driven by ITAM signaling pathways in macrophages is a known driver of autoimmune and inflammatory disease, with activated macrophages accumulating in chronically inflamed tissues in the absence of an active infection (68). Regulation of LynA protein expression and deactivation kinetics by c-Cbl, could, via the LynA checkpoint, help to prevent the initiation of pathological signaling. The threshold for macrophage activation is modifiable by changes to their local environment. For example, IFN-γ and LPS polarize macrophages for a pro-inflammatory response, whereas IL-4 and IL-13 polarize macrophages for tissue repair (i.e. collagen deposition) and inflammatory resolution (69, 70). We have reported that IFN-γ decreases the macrophage signaling threshold in part by increasing the expression of LynA (20). Dynamic changes in SFK and c-Cbl levels could modulate the macrophage activation threshold, ensuring that macrophages respond appropriately during times of infection (low threshold) and limiting aberrant activation in response to cellular debris and small-scale antibody complexes during inflammatory resolution (high threshold).

Like macrophages, mast cells reside in nearly every bodily tissue and perform environment-specific functions in addition to sensing non-self (69, 71). While macrophages have distinct anti-inflammatory roles as professional phagocytes in the silent clearance of apoptotic cells and agents of wound healing, mast cells are constitutively primed for ITAM-induced triggering, releasing preformed granules that contain inflammatory cytokines, chemokines, prostaglandins, and proteases. Pursuant to these differing functions, macrophages continuously gauge ITAM ligand valency (*i.e.* particle size) and have a relatively high basal threshold for inflammatory activation (5, 20), while mast cells can be triggered by small-scale receptor clustering induced by low-valency or even monovalent FcεR-IgE-allergen complexes (21, 22). One striking difference between the macrophage and mast-cell ITAM regulatory machinery is that mast cells express almost no c-Cbl. In spite of low reported mRNA levels (ImmGen, 51, 52), mast cells constitutively express a high level of LynA protein, likely due to impaired steady-state degradation of basally active LynA. Furthermore, LynA is not degraded upon activation and 3-IB-PP1 triggers a stronger Erk phosphorylation response in the absence of ITAM-receptor ligation in resting mast cells than in resting macrophages.

Overall, we describe a model in which regulation of LynA and c-Cbl levels helps to determine an immune cell’s potential for inflammatory ITAM signaling. Regulated degradation tunes down the LynA rheostat, blocking cell signaling via the LynA checkpoint at low cellular doses of LynA and overriding the LynA checkpoint at higher doses. In macrophages, this occurs on a continuum where LynA dose is highest in M1-polarized cells but still susceptible to rapid degradation by c-Cbl. Mast cells express almost no c-Cbl and maintain high levels of LynA at steady state and over time, consistent with permissive signaling in the absence of large-scale ITAM receptor clustering nucleated by multivalent receptor ligation. Appropriate regulation of immune receptor thresholds is critical for maintaining the function of innate immune cells; threshold dysregulation can lead to chronic feedback loops that drive inflammatory signaling in autoimmune disease and conversely tumor-supporting immunosuppressive signaling (68). Elucidating the mechanisms by which the LynA checkpoint is regulated may allow us to tune immune-cell sensitivity and reprogram pathological cells without causing immunodeficiency.

## Materials and Methods

### Mice

C57BL/6-derived *Csk^AS^* mice are hemizygous for the *Csk^AS^* BAC transgene on a *Csk^−/−^* background, as described previously (20, 26). Due to sterility of c-Cbl^−/−^ male mice, we were not able to produce a sustained lineage of *Csk^AS^Cbl^−/−^* mice, but we obtained three individuals by crossing *c-Cbl^−/−^* female mice from E. Peterson (University of Minnesota) (72) with *Csk^+/−^* (26) and *Csk^AS^* male mice from our colony and then crossing *Cbl^+/−^Csk^+/−^ with Cbl^+/−^Csk^AS^* mice. All mice were housed in specific pathogen-free conditions and genotyped using real-time PCR (Transnetyx, Inc., Memphis, TN).

### Jurkat cell lines and transfection

The Jurkat T-cell strains E6-1 (“wild-type”) (73) and JCaM1.6 (Lck-deficient) (43, 46) were gifts from the laboratories of A. Weiss (University of California, San Francisco) and Y. Shimizu (University of Minnesota), respectively. Jurkat cell lines were cultured in RPMI-1640 medium supplemented with 5% fetal calf serum (Omega Scientific, Inc., Tarzana, CA) and 2 mM glutamine, penicillin and streptomycin (Sigma-Aldrich, St. Louis, MO) as described previously *(74).* Jurkat and JCaM1.6 cells were transiently transfected via electroporation, as described previously (74). Briefly, cells were grown overnight in antibiotic-free RPMI-1640 medium supplemented with 10% fetal bovine serum (Omega Scientific) and 2 mM glutamine (RPMI10). Batches of 15 M cells were resuspended in RPMI10 with 15 µg plasmid DNA per construct. Cells were rested 20 min, electroporated 285 V/10 ms in a BTX square-wave electroporator (Harvard Apparatus, Holliston, MA), resuspended in 10 mL RPMI10, and allowed to recover overnight. One million live cells were then resuspended in phosphate-buffered saline, rested for 30 min at 37°C, and stimulated.

### DNA constructs and mutagenesis

Plasmids containing V5+His-tagged mouse LynA and LynB were gifts from J. Rivera/R. Suzuki (National Institutes of Health) (18). Myc-tagged mouse memCsk^AS^ (also referred to as Lck_11_-Csk^AS^) (25) and Xpress-tagged human c-Cbl and Cbl-b (75) were gifts from A. Weiss (University of California, San Francisco). Site-directed mutagenesis was performed using QuikChange Lightning (Agilent Technologies, Santa Clara, CA). Sequences of all constructs were verified by Sanger sequencing (GENEWIZ, South Plainfield, NJ).

### Preparation and flow cytometry of myeloid cells

BMDMs were prepared using standard methods (76). Briefly, bone marrow was extracted from femura and tibiae of mice aged 6-8-weeks. After hypotonic lysis of erythrocytes, BMDMs were derived on untreated plastic plates (BD Falcon, Sigma-Aldrich) by culturing in Dulbecco’s Modified Eagle Medium (DMEM, Corning Cellgro, Corning, NY) containing approximately 10% heat-inactivated fetal calf serum (Omega Scientific), 0.11 mg/ml sodium pyruvate (Corning), 2 mM penicillin/streptomycin/L-glutamine (Sigma-Aldrich), and 10% CMG-14-12-cell-conditioned medium as a source of M-CSF (77). After 6 or 7 days cells were resuspended in enzyme-free EDTA buffer and replated in untreated 6-well plates (BD Falcon, Sigma-Aldrich) at 1 M cells per well in unconditioned medium with or without priming in 25 U IFN-γ (PeproTech, Rocky Hill, NJ). Bone-marrow-derived mast cells were prepared as described previously (55), as above except for the substitution of 10 ng/ml IL-3 (PeproTech) for CMG-14-12-conditoned medium. After 5 weeks, cells were subjected to flow-cytometry analysis (LSRFortessa, Becton Dickinson, Franklin Lakes, NJ) and found to be uniformly positive for FcεRIα (Mar-1, FITC-labeled, #134305) and c-Kit (2B8, APC-labeled, #105811), both from BioLegend (San Diego, CA). Cells were rested or primed overnight before stimulation.

### siRNA knockdown

The siRNA sequence for knockdown of mouse c-Cbl was adapted from human siRNA. Cbl-b siRNA was designed using the Integrated DNA Technologies double-stranded siRNA design tools (IDT, Skokie, IL). Control double-stranded RNA from IDT was not predicted to be complementary to any sequence in human, mouse, or rat transcriptomes. 2 M BMDMs in 100 μL opti-MEM (ThermoFisher, Waltham, MA) were transfected with 1 μM siRNA via electroporation at 400 V/10 ms using a BTX square-wave electroporator. Cells were then plated on 150 mm untreated cell culture dishes and rested in 10 mL DMEM-10 for 30 min before adding 10 mL DMEM-10 supplemented with 10% CMG-14-12-cell-conditioned medium as a source of M-CSF. Transfections were pooled from several cuvettes to obtain enough cells for several stimulation conditions. After 24 h cells were resuspended in enzyme-free EDTA buffer and replated in untreated 6-well plates (BD Falcon, Sigma-Aldrich) at 1 M cells per well in unconditioned medium. Cells were used 48 h after transfection.

### Cell stimulation

BMDM stimulations have been described previously (20). After resting as described above, 1 M live Jurkat cells, mast cells, or adherent BMDMs were treated at 37°C in DMEM (myeloid cells) or PBS (Jurkat cells) with 10 µM 3-IB-PP1, a gift from K. Shokat (University of California, San Francisco). Signaling reactions were quenched by placing on ice and lysing cells in sodium dodecyl sulfate (SDS) buffer (128 mM Tris base, 10% glycerol, 4% SDS, 50 mM dithiothreitol, pH 6.8). Whole-cell lysates were prepared for immunoblotting by sonicating with a Bioruptor (Diagenode, Inc., Denville, NJ) for 3 min and boiling for 15 min.

### Antibodies and immunoblotting

After sample preparation, 0.25 M cell equivalents were run on each lane of a 7% NuPage Tris-Acetate gel (Invitrogen, Carlsbad, CA) and then transferred to an Immobilon-FL PVDF membrane (EMD Millipore, Burlington, MA). REVERT Total Protein Stain was used according to the standard protocol to quantify lane loading. After destaining, membranes were then treated with Odyssey® Blocking Buffer (TBS) for at least 1 h. Blotting was performed using standard procedures, and blots were imaged on an Odyssey CLx near-infrared imager (LI-COR Biosciences, Lincoln, NE). Antibodies for immunoblotting were purchased from Cell Signaling Technology (Danvers, MA), ProMab Biotechnologies (Richmond, CA), Thermo Fisher Scientific (Waltham, MA), LifeSpan Biosciences (Seattle, WA), Abcam (Cambridge, UK), and LI-COR Biosciences as follows:

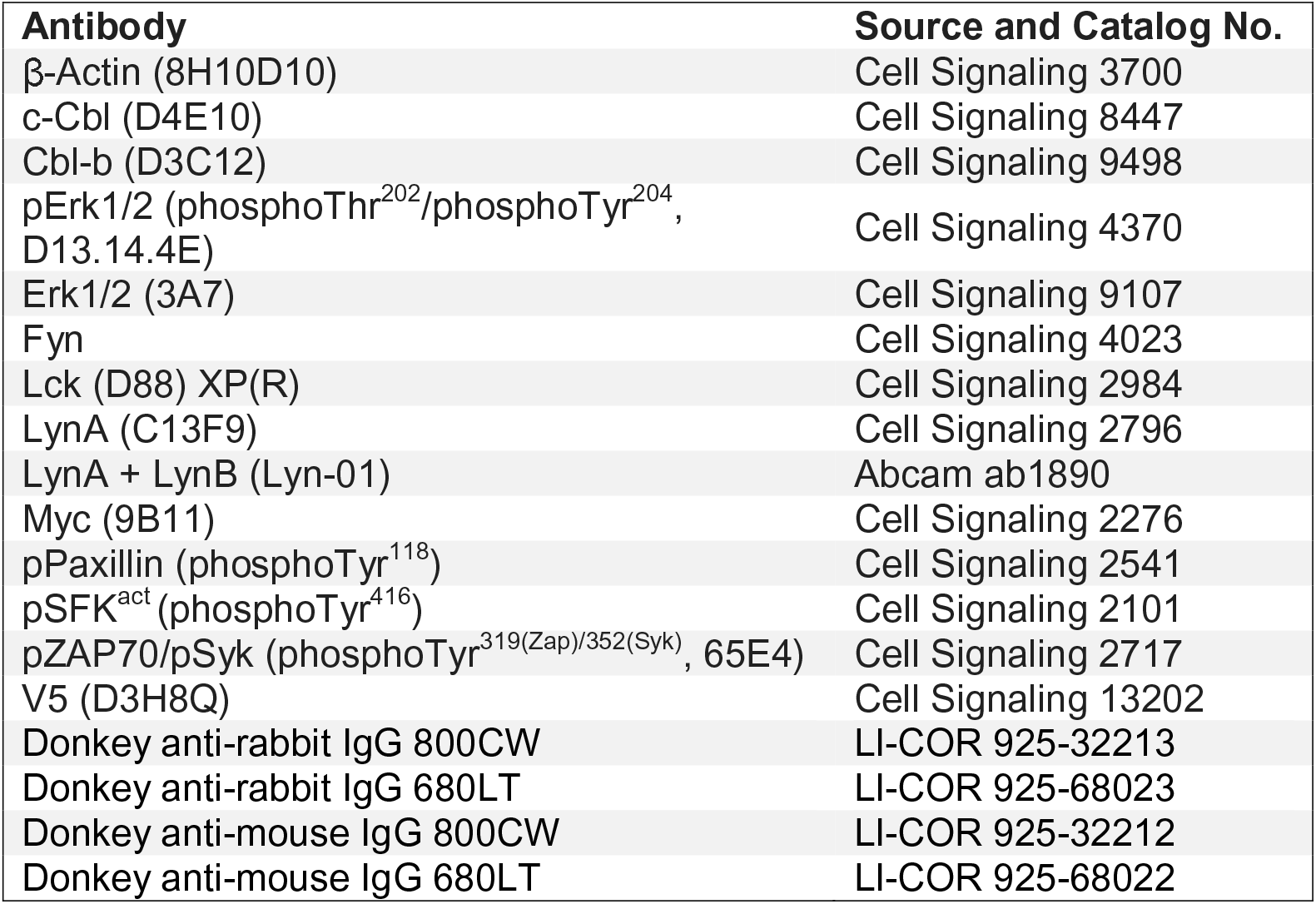

### Quantification, statistics, and image processing

Immunoblots were analyzed by densitometry from ImageStudio software (LI-COR Biosciences). Images were background-subtracted, and bands of the appropriate molecular weight were demarcated and analyzed for each gel lane. Each value was corrected for the total lane protein content (REVERT stain, above). Data were further normalized to a control condition to show relative changes in protein content. Where phosphoproteins were quantified, these were corrected for the total amount of the individual protein, rather than whole-lane protein content. For all figures, n values are biological replicates, reflecting independent experiments from different batches of cells, stimulus preparations, and experimental workflows. Statistical analyses were performed using Prism Software (Graphpad, La Jolla, CA). Significance was assessed using one- or two-way ANOVA analysis with Tukey’s or Sidak correction for multiple comparisons as indicated in the figure legends, with analysis of the mean at each time point or condition. Error bars in each figure represent standard errors of the mean from at least three independent experiments. Asterisks reflect specified P-values. Figures were further prepared using Adobe Creative Cloud software (San Jose, CA). Where appropriate, images were optimized by applying brightness/contrast changes to the whole image. No gamma correction was used. Images were only rotated after densitometry analysis was completed.

## Acknowledgements

We thank Arthur Weiss, Kevan Shokat, Yoji Shimizu, Brandon Burbach, Erik Peterson, Juan Rivera, and Ryo Suzuki for reagents and discussion and Terri Kadlecek, Marianne Mollenauer, and Whitney Swanson for technical support. Many thanks also to Hai-Bin Ruan, John Connett, Marc Jenkins, Bryce Binstadt, Erik Peterson, Nicholas Levinson, Aditi Bapat, Frances Sjaastad, and Emily Ewan for valuable discussions and feedback.

## Author Contributions

Conceptualization, Methodology, and Data Curation: BFB, TSF

Investigation, Validation, and Visualization: BFB, MGN, TSF

Formal Analysis, Writing- Review and Editing: BFB, MGN, KLS, TSF

Writing- Original Draft: BFB, TSF

Funding Acquisition: BFB, KLS, TSF

Project Administration, Resources, Validation, and Software: TSF

Supervision: KLS, TSF

## Data and Materials Availability

Reagents and protocols will be provided upon request.

## Figure Supplements

Figure 1–figure supplement 1. Loss of positive signaling in c-Cbl-deficient cells prevents assessment of LynA contributions to Erk phosphorylation.

Figure 3–figure supplement 1. LynA^Y32A^ and LynB are expressed more highly in Jurkat cells than wild-type LynA.

## Source Data

Figure 1–Source Data 1

Figure 3–Source Data 1

Figure 3–figure supplement 1-Source Data 1

Figure 4–Source Data 1

Figure 5–Source Data 1

Figure 6–Source Data 1

